# CTL1 (SLC44A1) regulates myelin lipid composition and architecture in Schwann cells

**DOI:** 10.64898/2026.09.24.753911

**Authors:** Akash A Patel, Corey Heffernan, Shakshi Desai, Andrew S. Lee, Nisha Gautam, Edward M. Bonder, Patrice Maurel, Haesun A. Kim

## Abstract

Myelin formation requires extensive membrane lipid synthesis, yet how myelinating glia acquire the choline needed for this process remains incompletely understood. Choline transporter-like protein 1 (CTL1; SLC44A1) is highly expressed in Schwann cells, and CTL1 deficiency in oligodendrocytes impairs CNS myelination. Here, we examined the role of CTL1 in peripheral nerve myelination using Schwann cell-specific *Ctl1* knockout mice. Unexpectedly, loss of *Ctl1* did not alter Schwann cell differentiation, the number of myelinated axons, or myelin thickness, but increased myelin abnormalities, including infoldings and outfoldings. Lipidomic analysis revealed selective alterations in myelin lipid composition, particularly among long-chain lipid species, together with triglyceride accumulation in whole nerves. CTL1 deficiency also increased mTORC1-associated S6 and mTORC2-associated AKT S473 phosphorylation. Transcriptomic analysis revealed downregulation of gene programs associated with fatty acid β-oxidation, triglyceride catabolism, and oxidative phosphorylation. Following peripheral nerve injury, *Ctl1*-deficient Schwann cells generated a normal repair response and efficiently remyelinated regenerated axons despite altered mTOR signaling. Together, these findings demonstrate that CTL1 is not required for overall peripheral myelin production but contributes to maintaining normal myelin lipid composition and architecture, revealing an unexpected ability of Schwann cells to sustain myelination despite disruption of CTL1-dependent choline metabolism

## INTRODUCTION

In both the peripheral (PNS) and central nervous system (CNS), glial cells generate a spiral wrapping of cell membrane around axons known as myelin. The myelin membrane is a specialized cellular membrane that not only allows for saltatory conduction but also plays additional roles in regulating axon caliber, ion channel organization, and providing trophic support to neurons (Eichel et al., 2020; Funfschilling et al., 2012; Lee et al., 2012; Salzer, 2003; Yin et al., 1998). This highly specialized membrane has a unique composition enriched with lipids, which make up 78% of the sheath compared to non-myelin tissue which contains only 35-40% lipid content (O’Brien & Sampson, 1965). Experimental deletion of many genes involved in lipid biosynthesis causes myelin defects, highlighting the requirement of lipid composition in myelin integrity (Schmitt, Castelvetri, & Simons, 2015).

Despite being generated by different glia in the PNS and CNS (Schwann cells and oligodendrocytes, respectively), myelin lipid compositions are quite similar. One notable difference is the greater representation of phospholipids and sphingomyelin in PNS myelin, which together account for approximately 42% of PNS myelin lipids compared with approximately 32% in the CNS (O’Brien, Sampson, & Stern, 1967; Poitelon, Kopec, & Belin, 2020). Phosphatidylcholine is a major phospholipid of cellular and myelin membranes, and its synthesis is closely dependent on cellular choline availability. Because mammalian cells cannot synthesize sufficient choline to meet their requirements, extracellular choline must be transported across the plasma membrane and incorporated into PC predominantly through the Kennedy pathway (Kenny, Scharenberg, Abu-Remaileh, & Birsoy, 2025). Indeed, approximately 70% of PC synthesis is estimated to derive from exogenous choline (Reo, Adinehzadeh, & Foy, 2002). Choline transport therefore represents a potentially important control point linking extracellular nutrient availability to membrane lipid synthesis in myelinating glia.

Choline transporter-like protein 1 (CTL1), encoded by *Slc44a1*, is a major intermediate-affinity choline transporter and is highly expressed throughout the nervous system (Traiffort, Ruat, O’Regan, & Meunier, 2005). Conditional deletion of *Slc44a1* from the oligodendrocyte lineage disrupts oligodendrocyte differentiation and reduces CNS myelination, demonstrating that adequate choline transport is critical for oligodendrocyte development and myelin production (Chen et al., 2025; Liu et al., 2025). However, whether Schwann cells have a comparable dependence on CTL1 for PNS myelination remains unknown.

Several observations implicate CTL1 in Schwann cells. A recent single-cell RNA-sequencing study indicates that *Ctl1* is the most highly expressed member of the *Ctl*-family in Schwann cells during development and into adulthood (Gerber et al., 2021). We previously showed that CTL1 is highly expressed in myelinating Schwann cells and is enriched along the axon-glia interface of the internode and in Schmidt-Lanterman incisures (Heffernan et al., 2017). Moreover, knockdown of *Ctl1* in cultured Schwann cells reduced myelin segment formation (Heffernan et al., 2017) suggesting that CTL1-mediated choline transport may contribute to Schwann cell myelination. These findings led us to hypothesize that CTL1 is required for PNS myelin formation and lipid homeostasis in vivo.

To test this hypothesis, we generated a mouse line with Schwann cell-specific deletion of CTL1 using Dhh-Cre (Dhh^Cre+^:*Ctl1^flox/flox^*) (*Ctl1* SC-KO). Unexpectedly, in contrast to the pronounced impairment of CNS myelination following CTL1 loss in oligodendrocytes, Schwann cells lacking CTL1 remained capable of generating myelin. However, *Ctl1* SC-KO nerves exhibited increased structural myelin abnormalities including infoldings and outfoldings. Loss of CTL1 also increased mTORC1 and mTORC2 signaling during Schwann cell development and following nerve injury, although these signaling changes did not produce a major alteration in remyelination. Lipidomic analysis further revealed accumulation of triglycerides and broader alterations in nerve lipid composition, indicating that CTL1 deficiency affects not only membrane lipid synthesis but also the metabolic handling and storage of lipids. Consistent with this interpretation, transcriptomic pathway analysis identified coordinated suppression of fatty-acid β-oxidation, triglyceride catabolism, and oxidative phosphorylation in *Ctl1* SC-KO nerves.

## Results

### Ctl1 is the major choline transporter expressed in Schwann cells

Recent studies have demonstrated that *Ctl1* is a major regulator of myelination in oligodendrocytes (Chen et al., 2025; Liu et al., 2025). *Ctl1* is a member of the *Ctl*-family of transporters which includes five intermediate-affinity transporters (*Ctl1*-*5*). One oligodendrocyte study showed that while *Ctl1* is the major choline transporter for myelination, other family members can also contribute to myelination (Liu et al., 2025). Analysis of mouse sciatic nerve transcriptomics indicates that *Ctl1* is the most highly expressed choline transporter in Schwann cells throughout development and into maturity, though other *Ctl*-family genes may also be expressed (Gerber et al., 2021). Therefore, we set out to determine which *Ctl*-family genes are expressed in Schwann cells. We generated cDNA from isolated rat Schwann cells and dorsal root ganglion (DRG) neuron cultures and analyzed the expression of *Ctl1-5*. cDNA from rat colon, which is known to express all *Ctl*-family members, was used as a positive control (Damm et al., 2017; O’Regan et al., 2000). All five *Ctl* genes were detected in DRG neuron cultures. In contrast, only *Ctl1* was detected in isolated Schwann cells (Figure 1A), indicating that *Ctl1* is the sole *Ctl*-family member expressed in Schwann cells. We next examined whether CTL1 affects intracellular choline levels in Schwann cells. We used cultured rat Schwann cells with shRNA-mediated *Ctl1* knockdown (sh-*Ctl1*), which we previously showed impairs myelin segment formation (Heffernan et al., 2017), with luciferase shRNA (sh-*Luc*) as the control. MALDI-TOF analysis showed a significant reduction in endogenous choline following *Ctl1* knockdown (Figure 1B), indicating that CTL1 is important for maintaining intracellular choline levels in Schwann cells.

**Figure 1.**
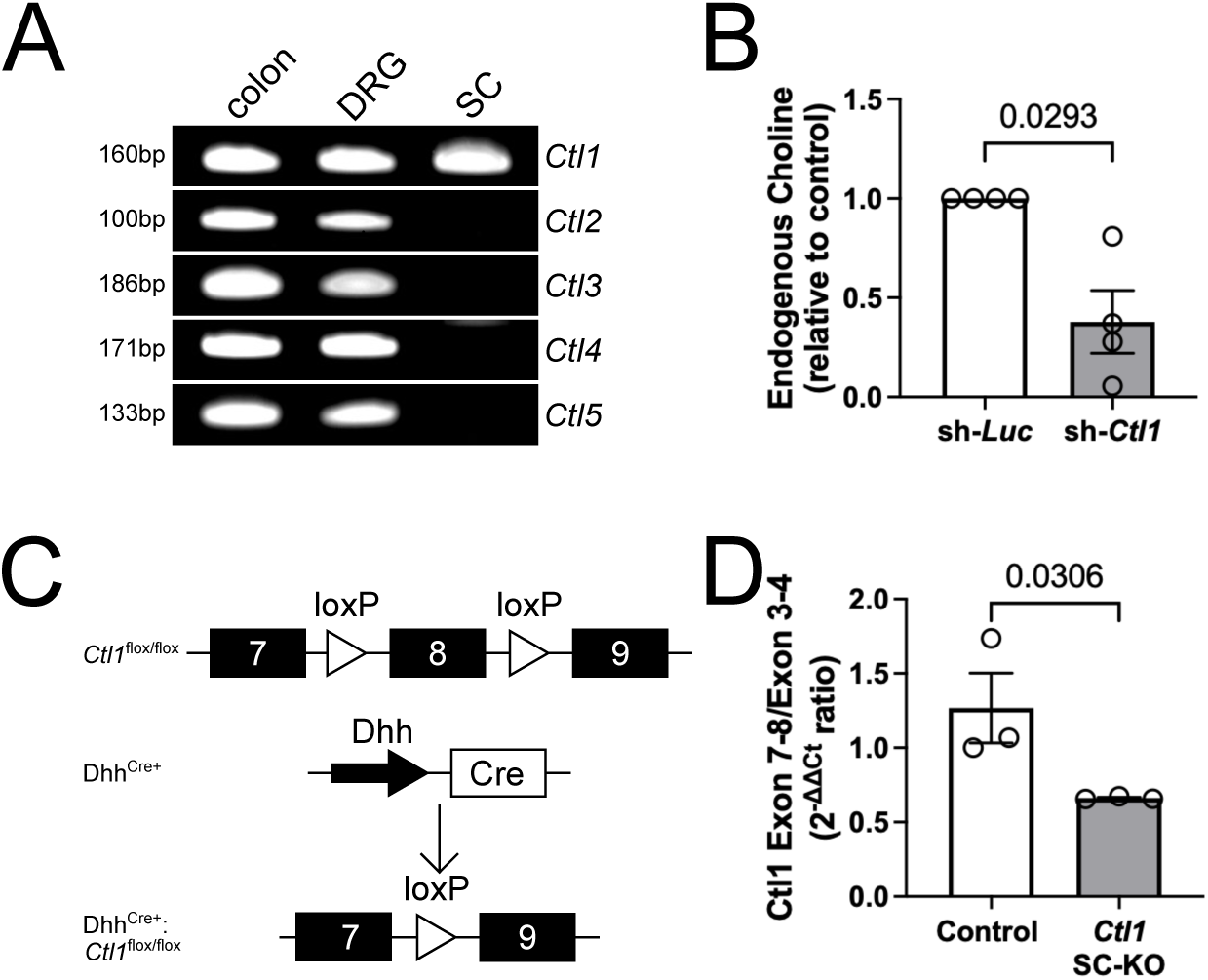
*Ctl1* is the major choline transporter for Schwann cells. **(A)** Image of RT-PCR products for *Ctl*-family genes for colon tissue, isolated dorsal root ganglion neurons (DRG), and isolated Schwann cells (SC). **(B)** Endogenous choline in sh-*Ctl1* Schwann cells relative to sh-*Luc* Schwann cells was assessed by MALDI-TOF mass spectrometry. n = 4. p = 0.0293. t(3) = 3.932. Two-tailed Paired t-test. **(C)** Diagram of *Ctl1* SC-KO model. **(D)** Graph of qPCR data measuring the ratio of products generated from primers of *Ctl1* spanning exons 7 and 8 relative to primers spanning exons 3 and 4. n = 3. p = 0.0306. t(4) = 2.582. One-tailed Unpaired t-test.

We next examined the role of *Ctl1* in Schwann cells, *in vivo*, by generating a Schwann cell-specific knockout mouse line (Figure 1C). The conditional *Ctl1* allele contains loxP sites flanking exon 8. Expression of Cre recombinase under the Schwann cell-specific *Dhh* promoter results in excision of exon 8, introducing a premature stop codon within the *Ctl1* open reading frame. To confirm exon 8 deletion, we generated cDNA from adult sciatic nerves of control and knockout mice and performed qPCR using primers spanning exons 7-8, with primers spanning exon 3-4 used to measure total *Ctl1* transcripts (Figure 1D). Nerves from the knockout mice showed a significant reduction in the relative abundance of exon 7-8 containing transcripts, consistent with efficient deletion of exon 8 in Schwann cells. Because cDNA was generated from whole sciatic nerves, the residual exon 7-8 signal in knockout nerves likely reflects *Ctl1* expression in the non-Schwann cell population.

### Ctl1 ablation causes myelin abnormalities

Choline is an essential precursor for phospholipids synthesis and is therefore particularly important during myelination, which requires extensive membrane biogenesis. In Schwann cells, membrane surface area increases approximately 50-fold during myelination to generate the 72-94 membrane wraps found in mature myelin (Webster, 1971). Consistent with the high demand for phospholipid synthesis, recent studies have demonstrated that *Ctl1* ablation in oligodendrocytes leads to reduced mature oligodendrocyte numbers, fewer myelinated axons, and CNS hypomyelination (Chen et al., 2025; Liu et al., 2025). We therefore asked whether *Ctl1* is similarly required for Schwann cell development and myelination in the adult PNS.

We first examined whether loss of *Ctl1* affects Schwann cell number or acquisition of the myelinating phenotype. In P30 sciatic nerves, the proportion of Schwann cells, marked by SOX10 expression, within adult sciatic nerves was unchanged in *Ctl1* SC-KO mice (Figure 2A and 2B) indicating that CTL1 deletion does not reduce the Schwann cell population. We next examined expression of KROX20, a key transcription factor for Schwann cell differentiation and myelination (Topilko et al., 1994). KROX20 expression was comparable between control and *Ctl1* SC-KO Schwann cells (Figure 2A and 2C). Consistent with preserved KROX20 expression, Western blot analysis showed no significant differences in the abundance of the major myelin proteins, myelin protein zero (MPZ) or myelin basic protein (MBP), between control and *Ctl1* SC-KO nerves (Figure 2D-F).

**Figure 2.**
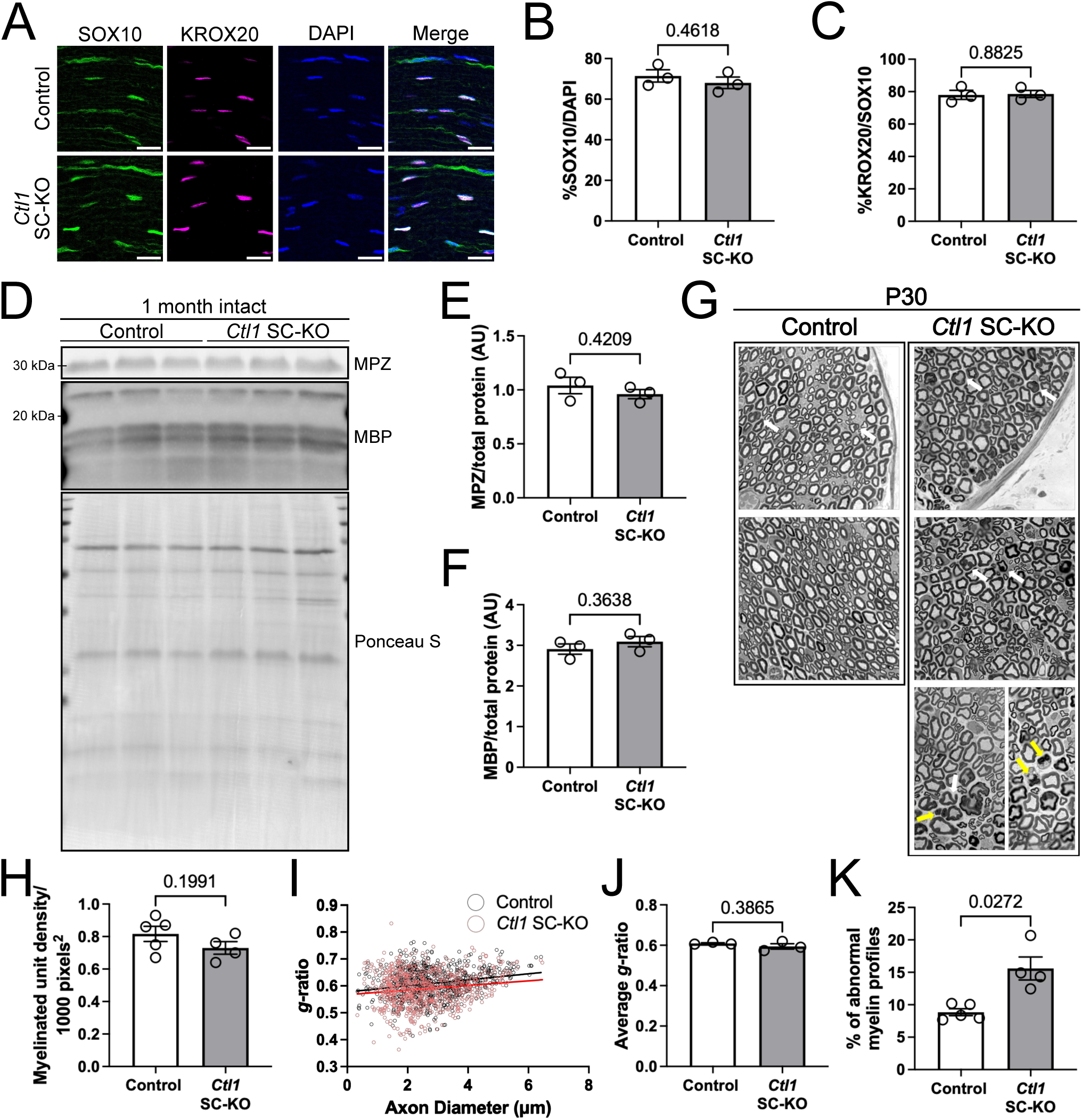
*Ctl1* regulates the generation of myelin abnormalities. **(A)** Representative immunostaining images of P30 control and *Ctl1* SC-KO nerves for SOX10 (green), KROX20 (magenta), and DAPI (blue). Scale bar = 20μm. **(B)** Graph of SOX10^+^ cell percentage. n = 3. p = 0.4618. t(3.9) = 0.8133. Two-tailed Unpaired t-test with Welch’s correction. **(C)** Graph of KROX20^+^ cell percentage in the SOX10^+^ population. n = 3. p = 0.8825. t(3.791) = 0.1580. Two-tailed Unpaired t-test with Welch’s correction. **(D)** Western blots of MPZ and MBP in one month control and *Ctl1* SC-KO intact sciatic nerves with Ponceau S-stained total protein loading control. **(E)** Graph of MPZ/total protein. n = 3. p = 0.4209. t(3.141) = 0.9240. Two-tailed Unpaired t-test with Welch’s correction. **(F)** Graph of MBP/total protein. n = 3. p = 0.3638. t(4) = 1.024. Two-tailed Unpaired t-test with Welch’s correction. **(G)** Representative toluidine blue-stained semithin images of control and *Ctl1* SC-KO nerves at P30. White arrows point to myelin abnormalities including infoldings and outfoldings, comma-shaped profiles, and irregularly shaped myelin–axon units. Yellow arrows point to degenerating myelin profiles. **(H)** Graph of myelinated axon-Schwann cell unit density across cross sections of control and *Ctl1* SC-KO nerves per 1000 square pixels. n = 4-5. p = 0.1991. t(6.988) = 1.418. Two-tailed Unpaired t-test with Welch’s correction. **(I)** Graph of *g*-ratios of intact sciatic nerves of control (black open circles) and *Ctl1* SC-KO (red open circles) of P30 mice. **(J)** Graph of average *g*-ratios control and *Ctl1* SC-KO. n = 3. p = 0.3865. t(2.22) = 1.071. Two-tailed Unpaired t-test with Welch’s correction. **(K)** Graph of the percentage of abnormal myelin profiles of intact sciatic nerves of control and *Ctl1* SC-KO of P30 mice. Abnormal myelin includes infoldings and outfoldings, comma-shapes, irregularly shaped myelin-axon units, and degenerating myelin. n = 4-5. p = 0.0272. t(3.539) = 3.634. Two-tailed Unpaired t-test with Welch’s correction.

In oligodendrocytes, deletion of *Ctl1* results in a reduced number of myelinated axons (Chen et al., 2025; Liu et al., 2025). We therefore first examined whether loss of *Ctl1* similarly affects peripheral nerve myelination. Morphological analysis was performed on semi-thin and ultra-thin sections of sciatic nerves from P30 control and *Ctl1* SC-KO mice (Figure 2G). There was no significant difference in the density of myelinated fibers between control and *Ctl1* SC-KO nerves (Figure 2H). Consistent with this finding, electron microscopy analysis of *g*-ratio shows that myelin thickness in *Ctl1* SC-KO is unchanged compared to control nerves (Figure 2I and 2J).

Despite the preservation of overall myelination, *Ctl1* SC-KO nerves showed a significant increase in abnormal myelin profiles compared with controls (Figure 2G and 2K). These abnormalities included myelin infoldings and outfoldings, comma-shaped myelin profiles, irregularly shaped myelin–axon units, and degenerating myelin. Thus, although loss of *Ctl1* does not substantially affect the number of myelinated axons or overall myelin thickness, it disrupts the normal architecture of the myelin sheath.

### Ctl1 ablation alters the long-chain lipid composition of peripheral myelin

Choline is an essential precursor for membrane lipid synthesis, and the specialized lipid composition of myelin is critical for its organization and function. Perturbations in lipid metabolism are associated with numerous peripheral neuropathies, and genetic disruption of lipid metabolic pathways can result in myelin abnormalities (Schmitt et al., 2015; Silva, Prior, D’Antonio, Swinnen, & Van Den Bosch, 2025). Recent lipidomic analysis of CNS myelin demonstrated that *Ctl1* deletion in oligodendrocytes causes substantial alterations in myelin lipid composition, including reductions in multiple phosphatidylcholine species (Chen et al., 2025). Given the increased myelin infoldings and outfoldings observed in *Ctl1* SC-KO nerves despite preservation of overall myelin thickness, we asked whether loss of *Ctl1* alters the lipid composition of peripheral myelin. We first performed untargeted lipidomic analysis of whole sciatic nerves. Surprisingly, *Ctl1* SC-KO nerves did not exhibit a broad depletion of lipids, with no lipid species decreased by more than two-fold relative to control nerves (Figure 3A). Instead, the most prominent changes were increases in several triglyceride species. Because triglycerides primarily represent storage lipids, their accumulation suggests altered lipid processing in *Ctl1*-deficient nerves but does not directly explain the abnormalities in myelin architecture.

**Figure 3.**
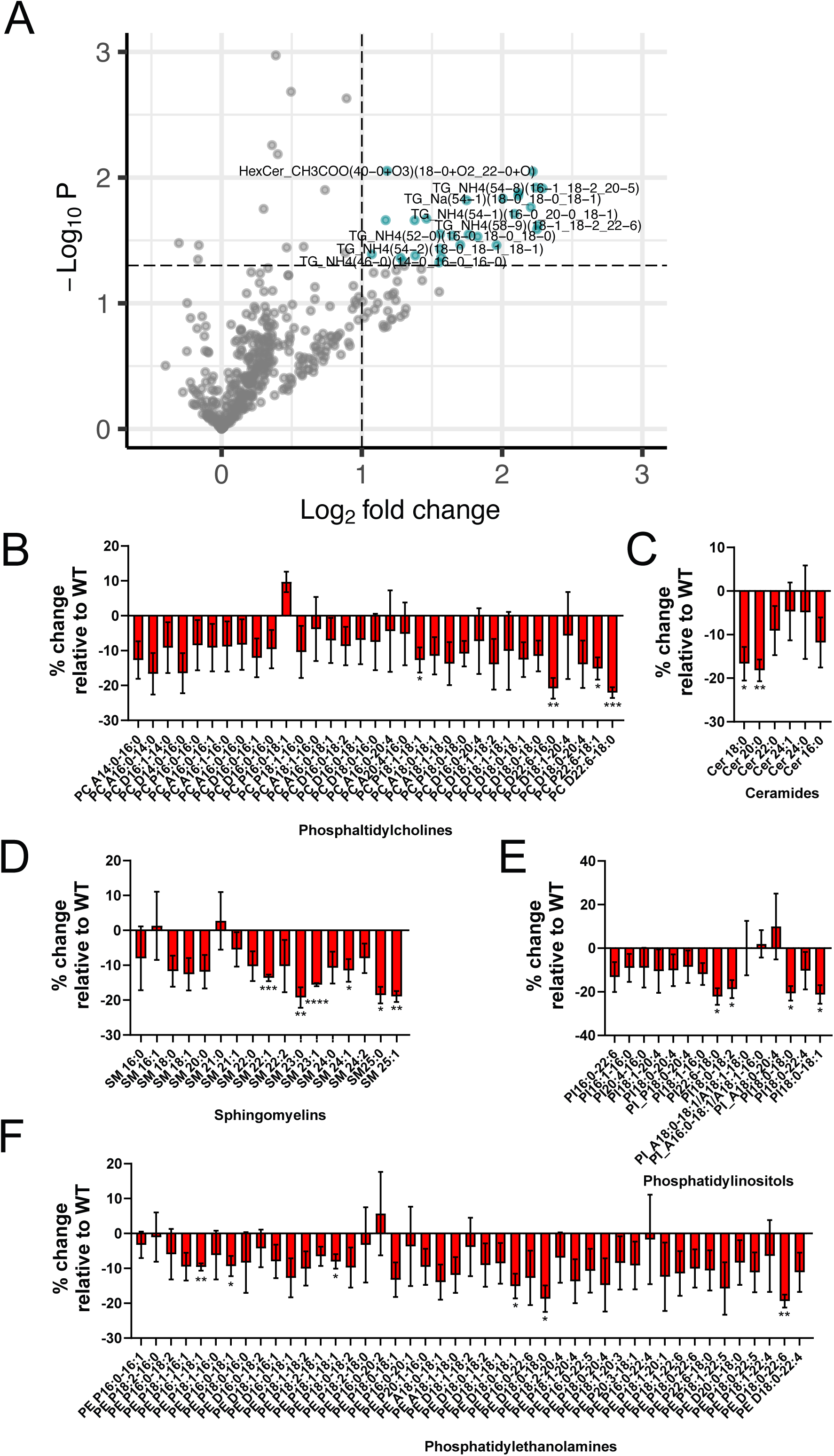
*Ctl1* ablation alters Schwann cell lipid content. (A) Volcano plots of differentially expressed lipids from whole sciatic nerves of control and *Ctl1* SC-KO. Log_2_FC ≥ |1| and p ≤ 0.05 represented as green circles; all others represented as gray circles. (B-F) Graph of abundance of (B) phosphatidylcholines, (C) ceramides, (D) sphingomyelins, (E) phosphatidylinositols, and (F) phosphatidylethanolamines in *Ctl1* SC-KO relative to wildtype control mice. No symbol, p > 0.05. *, p ≤ 0.05. **, p ≤ 0.01. ***, p ≤ 0.001. ****, p ≤ 0.0001. One sample t and Wilcoxon test.

We therefore examined the myelin membrane more directly by isolating a myelin-enriched fraction from sciatic nerves (Erwig et al., 2019) and performing targeted lipidomic analysis of major membrane lipid classes. Although most individual lipid species were not significantly altered, *Ctl1* SC-KO myelin showed selective reductions across several lipid classes, including phosphatidylcholines, ceramides, sphingomyelins, phosphatidylinositols, and phosphatidylethanolamines (Figure 3B–F). Notably, the reductions were enriched among lipid species containing long-chain and very-long-chain fatty acyl groups. Several phosphatidylcholine species containing C18–C22 fatty acids were decreased (Figure 3B), and prominent reductions were also observed among longer-chain sphingomyelin species (Figure 3D). Similarly, multiple phosphatidylinositol and phosphatidylethanolamine species containing C18–C22 fatty acyl chains were reduced in *Ctl1* SC-KO myelin (Figure 3E and 3F).

Thus, in contrast to the more extensive lipid depletion reported in *Ctl1*-deficient oligodendrocyte myelin (Chen et al., 2025), loss of *Ctl1* in Schwann cells produces a more selective remodeling of peripheral myelin lipids. Importantly, these changes preferentially affect lipid species containing longer fatty acyl chains rather than causing a generalized loss of myelin lipids.

### Ctl1 SC-KO nerves exhibit increased AKT–mTOR signaling

Myelin infoldings and outfoldings similar to those observed in *Ctl1* SC-KO nerves are characteristic of dysregulated PI3K–AKT–mTOR signaling. The PI3K–AKT–mTOR pathway plays a critical role in the initiation of Schwann cell myelination and regulation of myelin growth. Accordingly, constitutive activation of PI3K or AKT (Domenech-Estevez et al., 2016; Ishii, Furusho, & Bansal, 2021), as well as deletion of negative regulators of the pathway, including PTEN, TSC1, and TSC2 (Beirowski, Wong, Babetto, & Milbrandt, 2017; Figlia, Norrmen, Pereira, Gerber, & Suter, 2017; Goebbels et al., 2010), result in abnormal myelin growth characterized by myelin infoldings and outfoldings. Given the similar myelin abnormalities observed in *Ctl1* SC-KO nerves, we asked whether loss of *Ctl1* alters AKT–mTOR signaling in peripheral nerves.

We first examined AKT activation in sciatic nerves from P30 control and *Ctl1* SC-KO mice. AKT activation is regulated by phosphorylation at two major sites, T308 and S473. Phosphorylation of T308, which is mediated by PDK1 downstream of PI3K and is critical for AKT kinase activation, was unchanged in *Ctl1* SC-KO nerves (Figure 4A top and 4B). In contrast, phosphorylation of AKT at S473 was significantly increased in *Ctl1* SC-KO nerves (Figure 4A middle and 4C). Because AKT S473 is phosphorylated by mTORC2 (Manning & Toker, 2017), these findings suggest that loss of *Ctl1* preferentially enhances the mTORC2–AKT arm of the pathway rather than producing a generalized increase in upstream PI3K-dependent AKT activation.

**Figure 4.**
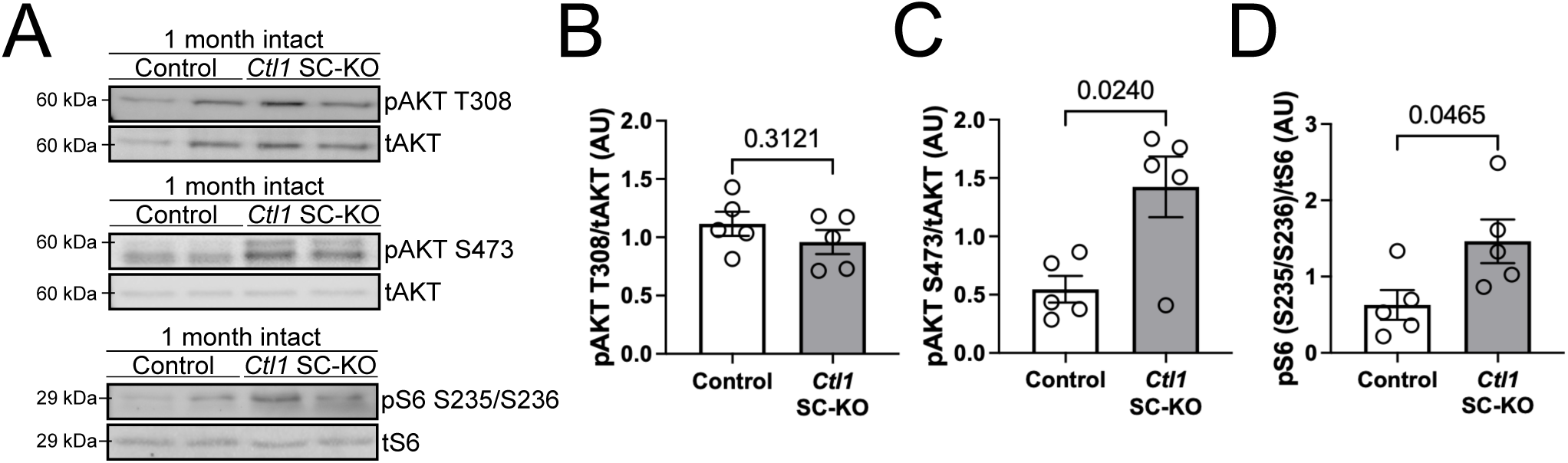
*Ctl1* ablation alters mTORC1 and mTORC2 signaling in intact nerves. (A) Representative Western blots of pAKT T308 and tAKT (top), pAKT S473 and tAKT (middle), and pS6 S235/236 and tS6 (bottom). (B) Graph of pAKT T308/tAKT. n = 5. p = 0.3121. t(8) = 1.079. Two-tailed Unpaired t-test with Welch’s correction. (C) Graph of pAKT S473/tAKT. n = 5. p = 0.0240. t(5.490) = 3.092. Two-tailed Unpaired t-test with Welch’s correction. (D) Graph of pS6 S235/236/tS6. n = 5. p = 0.0465. t(7.042) = 2.412. Two-tailed Unpaired t-test with Welch’s correction.

The increase in AKT S473 phosphorylation prompted us to determine whether mTORC1 signaling was also elevated. We therefore examined phosphorylation of ribosomal protein S6 at S235/S236, a downstream readout of mTORC1 kinase signaling (Biever, Valjent, & Puighermanal, 2015). S6 S235/S236 phosphorylation was significantly increased in *Ctl1* SC-KO sciatic nerves compared with controls (Figure 4A bottom and 4D).

Together, the selective increase in AKT S473 phosphorylation and increased phosphorylation of S6 indicate enhanced AKT–mTOR signaling in *Ctl1* SC-KO nerves. The absence of increased AKT T308 phosphorylation argues against broad activation of the canonical PI3K–PDK1–AKT pathway and instead suggests altered regulation of mTOR signaling downstream of, or independently from, proximal PI3K activation.

### Loss of Ctl1 alters transcriptional programs associated with lipid and oxidative metabolism

Given the alterations in lipid composition and mTOR signaling observed in *Ctl1* SC-KO nerves, we next asked whether loss of CTL1 alters metabolic gene programs in adult peripheral nerves. Bulk RNA-seq was performed on sciatic nerves from adult control and *Ctl1* SC-KO mice. Surprisingly, despite the pronounced lipid abnormalities described above, loss of *Ctl1* produced relatively few large changes in individual gene expression. Consistent with efficient deletion, *Slc44a1* (*Ctl1*) was among the most strongly downregulated genes, together with *Glp1r, Hal and Acpp*, whereas *Atp8b5* was increased (Figure 5A). Thus, the metabolic phenotype of *Ctl1* deficiency does not appear to result from broad transcriptional disruption but instead suggested more coordinated changes across functionally related gene networks.

**Figure 5:**
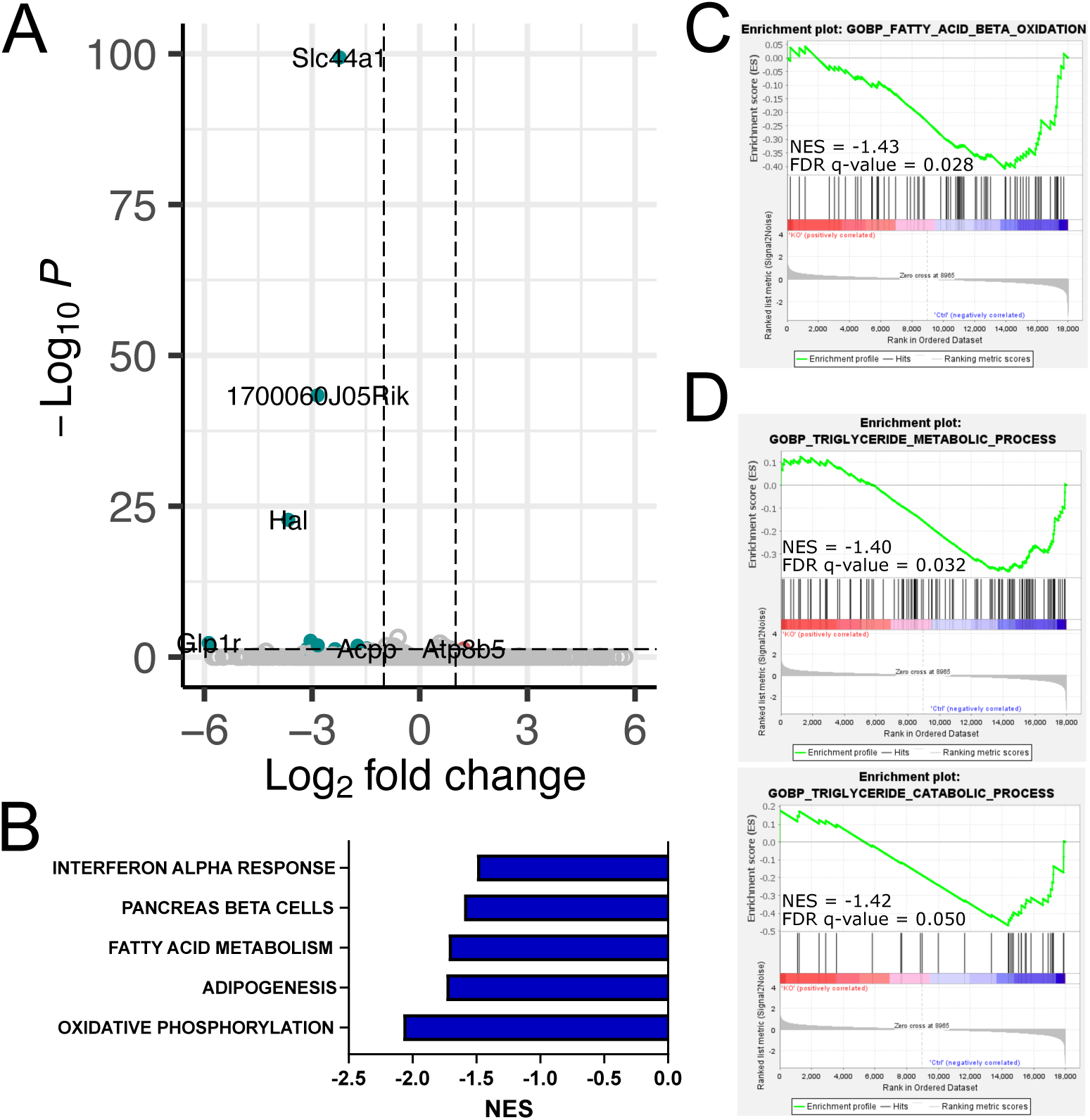
*Ctl1* ablation alters transcriptional programs associated with lipid and oxidative metabolism. (A) Volcano plot of DEGs in *Ctl1* SC-KO vs control intact nerves. DEGs with Log_2_FC ≥ |1| and p_adj_ ≤ 0.05 represented as red circles, Log_2_FC ≤ |1| and p_adj_ ≤ 0.05 represented as green circles, and all others represented as gray circles. (B) Graph of significantly downregulated GSEA hallmark pathways in *Ctl1* SC-KO vs control. (C) GSEA enrichment plot of *Ctl1* SC-KO vs control for “GOBP Fatty Acid Beta Oxidation.” (D) GSEA enrichment plot of *Ctl1* SC-KO vs control for “GOBP Triglyceride Metabolic Process” (top) and “GOBP Triglyceride Catabolic Process” (bottom).

To identify such pathway-level alterations, we performed gene set enrichment analysis (GSEA) (Ben-Porath et al., 2008; Liberzon et al., 2015). Strikingly, oxidative phosphorylation was the most negatively enriched Hallmark gene set in *Ctl1* SC-KO nerves, with fatty acid metabolism also among the most strongly downregulated pathways (Figure 5B). These findings indicate that CTL1 deficiency is associated with the coordinated suppression of pathways involved in mitochondrial energy production and lipid utilization, despite relatively modest changes in individual transcripts.

Because fatty acids represent an important oxidative substrate and our lipidomic analyses demonstrated increased triglyceride accumulation in *Ctl1* SC-KO nerves, we examined lipid catabolic pathways in greater detail. GSEA analysis of gene ontology biological process sets revealed significant negative enrichment of fatty acid β-oxidation in *Ctl1* SC-KO nerves (Figure 5C). Moreover, both triglyceride metabolic process and triglyceride catabolic process gene sets were negatively enriched (Figure 5D). Together, these changes suggest impaired mobilization of stored triglycerides, coupled with a reduced capacity for mitochondrial fatty acid oxidation. Importantly, these transcriptional changes provide a potential mechanistic link to the increased triglyceride accumulation in *Ctl1* SC-KO nerves (Figure 3A). It is possible that loss of Ctl1 suppresses pathways required for oxidizing fatty acids, thus shifting Schwann cell lipid metabolism away from utilization and toward storage.

### Ctl1 loss enhances mTORC1 signaling after nerve injury without altering early repair Schwann cell responses

Following peripheral nerve injury, myelinating Schwann cells undergo extensive reprogramming to generate repair Schwann cells that support axon regeneration. This transition is initiated rapidly after injury and involves activation of multiple signaling pathways that coordinate Schwann cell dedifferentiation, proliferation, and acquisition of the repair phenotype (Babetto, Wong, & Beirowski, 2020; Guertin, Zhang, Mak, Alberta, & Kim, 2005; Norrmen et al., 2018). Both mTORC1 and mTORC2 signaling have been implicated in regulating the Schwann cell response to nerve injury (Daboussi et al., 2023; Norrmen et al., 2018). Because *Ctl1* SC-KO nerves exhibited elevated AKT–mTOR signaling under uninjured conditions, we asked whether loss of *Ctl1* alters the early Schwann cell response following nerve injury. Sciatic nerves were transected, and distal nerve segments were analyzed at 3 days post-injury (DPI), when Schwann cells are actively transitioning to the repair phenotype.

We first examined mTOR signaling specifically in Schwann cells. Co-immunostaining for phosphorylated S6 S235/S236 and SOX10 showed a significant increase in the percentage of pS6-positive Schwann cells in *Ctl1* SC-KO nerves compared with controls (62 ± 3.4% vs. 47 ± 4.0%; Figure 6A), indicating increased mTORC1-associated signaling during the early response to injury. In contrast, the percentage of Schwann cells positive for AKT S473 phosphorylation, a readout of mTORC2 activity, was similar between control and *Ctl1* SC-KO nerves (63 ± 12.2% vs. 64 ± 9.9%, respectively) (Figure 6B). Thus, although *Ctl1* SC-KO nerves exhibit increased AKT S473 and S6 S235/S236 phosphorylation under uninjured conditions, an increase in injury-induced AKT S473 phosphorylation was not evident in distal Schwann cells. Instead, S6 S235/S236 phosphorylation was enhanced, suggesting that loss of *Ctl1* selectively enhances mTORC1-associated signaling during the early Schwann cell injury response.

**Figure 6.**
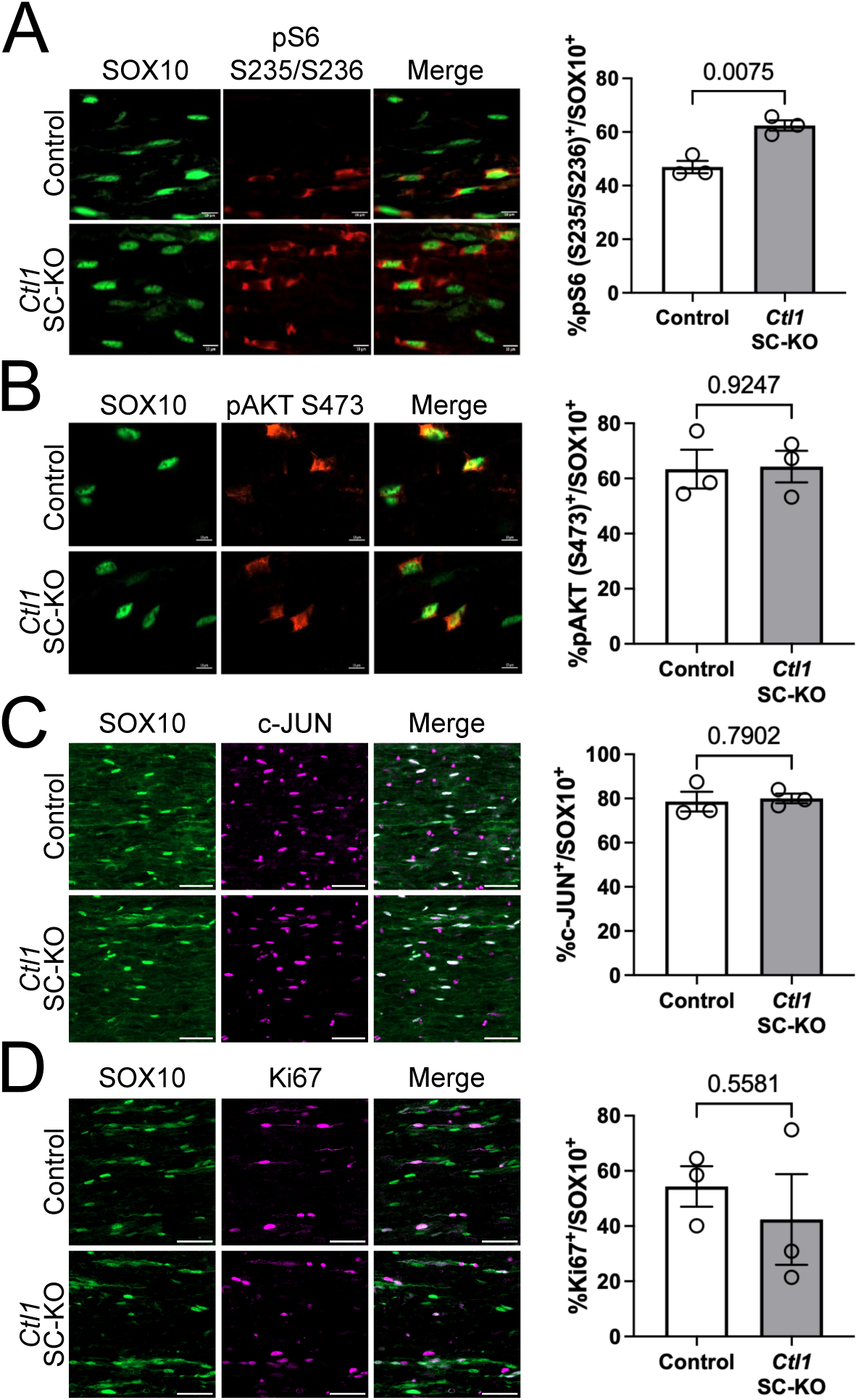
*Ctl1* ablation causes increased mTORC1 signaling early after injury without altering early repair response. **(A)** Representative immunostaining images of 3DPI transection control and *Ctl1* SC-KO nerves for SOX10 (green) and pS6 S235/S236 (red) (left). Scale bar = 10μm. Graph of pS6 S235/S236^+^ cell percentage in the SOX10^+^ population (right). n = 3. p = 0.0075. t(3.882) = 5.121. Two-tailed Unpaired t-test with Welch’s correction. (B) Representative immunostaining images of 3DPI transection control and *Ctl1* SC-KO nerves for SOX10 (green) and pAKT S473 (red) (left). Scale bar = 10μm. Graph of pAKT S473^+^ cell percentage in the SOX10^+^ population (right). n = 3. p = 0.9247. t(3.849) = 0.1009. Two-tailed Unpaired t-test with Welch’s correction. (C) Representative immunostaining images of 3DPI transection control and *Ctl1* SC-KO nerves for SOX10 (green) and c-JUN (magenta) (left). Scale bar = 20μm. Graph of c-JUN^+^ cell percentage in the SOX10^+^ population (right). n = 3. p = 0.7902. t(2.878) = 0.2919. Two-tailed Unpaired t-test with Welch’s correction. (D) Representative immunostaining images of 3DPI transection control and *Ctl1* SC-KO nerves for SOX10 (green) and Ki67 (magenta) (left). Scale bar = 20μm. Graph of Ki67^+^ cell percentage in the SOX10^+^ population (right). n = 3. p = 0.5581. t(2.763) = 0.6636. Two-tailed Unpaired t-test with Welch’s correction.

We next asked whether increased S6 phosphorylation was accompanied by altered acquisition of the repair Schwann cell phenotype. mTORC1 signaling contributes to regulation of c-JUN, a key transcription factor induced following nerve injury and required for major aspects of the repair Schwann cell program (Arthur-Farraj et al., 2012; Norrmen et al., 2018; Parkinson et al., 2008). Despite increased S6 phosphorylation, the percentage of c-JUN-positive Schwann cells was comparable between control (79 ± 7.8%) and *Ctl1* SC-KO (80 ± 3.7%) nerves at 3 DPI (Figure 6C). Thus, enhanced mTORC1-associated signaling in *Ctl1*-deficient Schwann cells does not result in increased or impaired induction of c-JUN during the early injury response.

Cell-cycle re-entry and proliferation are other prominent components of the early Schwann cell response to injury and can be regulated independently of c-JUN, but in part through mTORC1-dependent mechanisms (Abercrombie & Johnson, 1946; Arthur-Farraj et al., 2012; Norrmen et al., 2018; Stierli et al., 2018). We therefore examined Schwann cell proliferation by co-immunostaining for Ki67 and SOX10 at 3 DPI. The percentage of Ki67-positive Schwann cells did not differ significantly between control and *Ctl1* SC-KO nerves (Figure 6D).

Together, these results demonstrate that loss of *Ctl1* selectively enhances mTORC1-associated signaling during the early response to peripheral nerve injury without measurably altering c-JUN induction or Schwann cell proliferation. Thus, despite increased S6 phosphorylation, *Ctl1*-deficient Schwann cells retain the capacity to initiate key components of the early repair program. These findings further suggest that the elevated mTORC1 signaling caused by *Ctl1* loss is not sufficient to substantially alter early repair Schwann cell generation.

### Ctl1-deficient Schwann cells retain normal remyelinating capacity despite sustained mTOR hyperactivation

Following axon regeneration, repair Schwann cells transition from the repair state and re-establish a myelinating phenotype around regenerated axons (Stassart & Woodhoo, 2021). Because loss of *Ctl1* altered mTOR signaling in both intact nerves and during the early response to injury, we asked whether this signaling abnormality persists during the later phase of regeneration, when Schwann cells begin to remyelinate axons. We used a sciatic nerve crush injury model and collected distal nerves from control and *Ctl1* SC-KO mice at 18 DPI, a period of active remyelination (Patel et al., 2024). We first examined whether the altered mTOR signaling observed in *Ctl1* SC-KO nerves persisted at this stage. S6 S235/S236 phosphorylation was significantly increased in *Ctl1* SC-KO nerves (Figure 7A top and 7B), indicating sustained elevation of mTORC1-associated signaling. AKT S473 phosphorylation was also significantly increased (Figure 7A middle and 7C), consistent with increased mTORC2-associated signaling, whereas AKT T308 phosphorylation remained unchanged (Figure 7A bottom and 7D). Thus, similar to intact nerves, *Ctl1* SC-KO nerves during the remyelination phase exhibit increased mTORC1-and mTORC2-associated signaling without a corresponding increase in AKT T308 phosphorylation.

**Figure 7.**
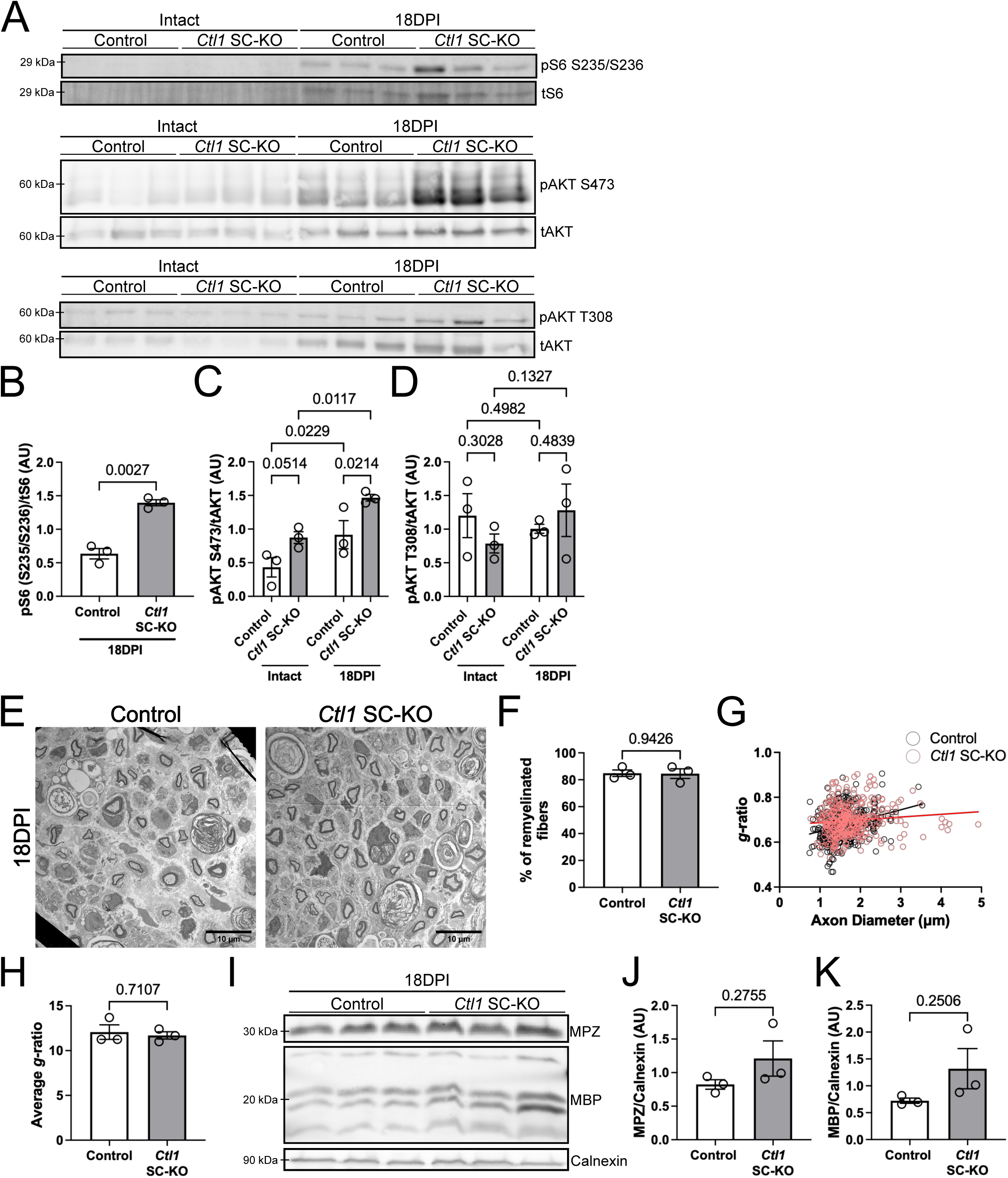
*Ctl1* absence-induced mTOR signaling does not alter remyelination. (A) Representative Western blots of contralateral intact and ipsilateral 18DPI control and *Ctl1* SC-KO for pS6 S235/236 and tS6 (top), pAKT S473 and tAKT (middle), and pAKT T308 and tAKT (bottom). (B) Graph of pS6 S235/236/tS6. n = 3. p = 0.0027. t(3.204) = 8.422. Two-tailed Unpaired t-test with Welch’s correction. (C) Graph of pAKT S473/tAKT. n = 3. p_Intact_ = 0.0514, p_18DPI_ = 0.0214, p_Control_ = 0.0229, and p_*Ctl1* SC-KO_ = 0.0117. t_Intact_(8) = 2.289, t_18DPI_(8) = 2.853, t_Control_(4) = 3.592, and t_*Ctl1* SC-KO_(4) = 4.399. 95% Confidence interval: CI_Intact_ = [-0.886, 0.003339], CI_18DPI_ = [-0.9947, -0.1054], CI_Control_ = [-0.8590, - 0.1101], and CI_*Ctl1* SC-KO_ = [-0.9677, -0.2188]. Two-way ANOVA with Uncorrected Fisher’s LSD. Only planned comparisons displayed within graph were analyzed. (D) Graph of pAKT T308/tAKT. n = 3. p_Intact_ = 0.3028, p_18DPI_ = 0.4839, p_Control_ = 0.4982, and p_*Ctl1* SC-KO_ = 0.1327. t_Intact_(8) = 1.101, t_18DPI_(8) = 0.7341, t_Control_(4) = 0.7441, and t_*Ctl1* SC-KO_(4) = 1.884. 95% Confidence interval: CI_Intact_ = [-0.4535, 1.283], CI_18DPI_ = [-1.144, 0.5918], CI_Control_ = [-0.5343, 0.9255], and CI_*Ctl1* SC-KO_ = [-1.225, 0.2346]. Two-way ANOVA with Uncorrected Fisher’s LSD. Only planned comparisons displayed within graph were analyzed. (E) Representative transmission electron micrographs of control and *Ctl1* SC-KO sciatic nerve cross sections at 18DPI. Scale bar = 10μm. (F) Graph of *g*-ratios of 18DPI sciatic nerves of control (black open circles) and *Ctl1* SC-KO (red open circles). (G) Graph of average *g*-ratios control and *Ctl1* SC-KO. n = 3. p = 0.7107. t(2.942) = 0.4087. Two-tailed Unpaired t-test with Welch’s correction. (H) Graph of percentage of remyelinated large axons (>1μm) of control and *Ctl1* SC-KO 18DPI nerves. n = 3. p = 0.9426. t(3.539) = 0.07722. Two-tailed Unpaired t-test with Welch’s correction. (I) Western blots of MPZ and MBP in 18DPI control and *Ctl1* SC-KO sciatic nerves with calnexin as loading control. (J) Graph of MPZ/Calnexin. n = 3. p = 0.2755. t(2.288) = 1.424. Two-tailed Unpaired t-test with Welch’s correction. (K) Graph of MBP/Calnexin. n = 3. p = 0.2506. t(2.067) = 1.582. Two-tailed Unpaired t-test with Welch’s correction.

We next determined whether this persistent increase in mTOR signaling alters Schwann cells’ ability to remyelinate regenerated axons. Because mTORC1 activity is required for timely remyelination and restoration of myelin thickness following peripheral nerve injury (Norrmen et al., 2018), we considered whether its sustained elevation in *Ctl1* SC-KO nerves might accelerate or otherwise alter remyelination. Ultrastructural analysis at 18 DPI revealed extensive remyelination in both control and *Ctl1* SC-KO nerves (Figure 7E). The percentage of remyelinated fibers was nearly identical between genotypes 85 ± 4.2% for the control and 85 ± 6.2% for the *Ctl1* SC-KO (Figure 7F), indicating that loss of *Ctl1* neither delays nor accelerates the onset of remyelination at this time point. Analysis of *g*-ratio revealed similar distributions between control and *Ctl1* SC-KO nerves (Figure 7G) with similar mean *g*-ratios between genotypes (Figure 7H). Consistent with the morphological analysis, expression of the major peripheral myelin proteins MPZ and MBP was also comparable between control and *Ctl1* SC-KO nerves at 18 DPI (Figure 7I-K).

Together, these findings demonstrate that *Ctl1*-deficient Schwann cells retain the capacity to efficiently remyelinate regenerated axons despite persistent dysregulation of mTOR signaling. Neither the extent of remyelination nor the major protein composition of newly formed myelin was detectably altered at 18DPI. These findings parallel the preservation of developmental myelination in *Ctl1* SC-KO nerves and indicate that CTL1 is dispensable for the bulk production of peripheral myelin during both development and regeneration. Importantly, they also demonstrate that sustained elevation of mTOR signaling in *Ctl1*-deficient nerves is not sufficient to increase the rate or extent of remyelination.

## Discussion

Choline is an essential nutrient that is required for the synthesis of major membrane lipids, including phosphatidylcholine and sphingomyelin, through the Kennedy pathway (Kennedy & Weiss, 1956; Kenny et al., 2025; Reo et al., 2002). Choline-derived lipids are major components of the myelin sheath in both PNS and CNS (Hendelman & Bunge, 1969; Poitelon et al., 2020). Recent studies in the CNS identified CTL1 as a critical choline transporter required for oligodendrocyte development and myelination (Chen et al., 2025; Liu et al., 2025). In contrast, our findings show that CTL1 plays a more limited role in Schwann cell myelination. Whereas deletion of *Ctl1* in oligodendrocytes reduces mature oligodendrocyte numbers, the number of myelinated axons, and myelin thickness (Chen et al., 2025; Liu et al., 2025), CTL1-deficient Schwann cells were able to generate myelin. Large axons were normally myelinated, and overall myelin thickness was not altered. However, the absence of *Ctl1* increased structural abnormalities of the myelin sheath, including infoldings and outfoldings. Thus, CTL1 is not essential for Schwann cells to generate myelin but is important for maintaining normal myelin architecture.

The relatively mild myelination phenotype was unexpected given the requirement for CTL1 in oligodendrocytes, and our previous finding that *Ctl1* knockdown impairs myelin formation in Schwann cell cultures. Furthermore, *Ctl1*-knockdown Schwann cells exhibit reduced endogenous choline levels. One possibility is that Schwann cells have alternative mechanisms for maintaining choline availability in vivo. In our myelin lipidomics, *Ctl1* SC-KO nerves showed reduced levels of selected phosphatidylcholines, ceramides, sphingomyelins, phosphatidylinositols, and phosphatidylethanolamines, consistent with altered myelin lipid composition. Nevertheless, Schwann cells were able to produce sufficient membrane to generate myelin sheaths. While, imported choline accounts for a large proportion of choline-derived lipids, choline can also be generated through the PEMT pathway (Reo et al., 2002). In Schwann cells, PEMT, is expressed during early postnatal development but becomes undetectable by P14 (Heffernan et al., 2017). However, disruption of choline transport in cultured Schwann cells increased lipids potentially derived from the PEMT pathway (Heffernan et al., 2017), and increased PEMT expression can compensate for choline deprivation in other cell types (Cui & Vance, 1996). Schwann cells may therefore increase endogenous choline production when CTL1 dependent transport is reduced.

Alternatively, other choline transporters may compensate for CTL1 *in vivo*. While not detected in cultured Schwann cells, previous transcriptomic studies show very low expression of *Ctl3*, *Ctl4*, and *Ctl5* in developing peripheral nerves, whereas *Ctl1* and *Ctl2* expression increases during postnatal nerve development into adulthood (Gerber et al., 2021). CTL1 and CTL2 both function as choline transporters and localize to the cell membrane and mitochondria (Taylor, Grapentine, Ichhpuniani, & Bakovic, 2021). CTL2 could therefore provide an alternative source of choline in Schwann cells in vivo. The striking difference between Schwann cells and oligodendrocytes suggests that the two myelinating glial cell types differ in how they acquire or maintain the choline needed for membrane synthesis. Defining these compensatory mechanisms will be important for understanding why Schwann cells remain capable of myelination in the absence of CTL1.

Although overall myelin formation was preserved, the lipid abnormalities indicate that compensation for CTL1 loss is incomplete. Myelin structure depends not only on the amount of membrane produced but also on its lipid composition (Poitelon et al., 2020; Schmitt et al., 2015; Silva, Prior, D’Antonio, Swinnen, & Van Den Bosch, 2025). Thus, Schwann cells may generate sufficient membrane to form a myelin sheath while failing to maintain the lipid composition required for normal myelin architecture. This may explain why *Ctl1* SC-KO nerves show relatively normal myelin thickness and numbers of myelinated axons while exhibiting increased abnormalities.

Our findings also suggest a broader relationship between CTL1 and lipid metabolism. Whole-nerve lipidomics did not reveal an overall depletion of choline-derived lipids in *Ctl1* SC-KO nerves but instead showed increased levels of several triglyceride species. Transcriptomic analysis similarly revealed changes in metabolic gene programs, including reduced expression of pathways associated with oxidative phosphorylation and fatty acid β-oxidation. Similar relationships between impaired choline metabolism, triglyceride accumulation, and altered mitochondrial metabolism have been reported in other systems (Fagerberg et al., 2020; Lu et al., 2023; Michel, Singh, & Bakovic, 2011; Taylor, Schenkel, Yokich, & Bakovic, 2017). Fatty acid oxidation flux, mitochondrial respiration, and ATP production were not directly measured in our study. Nevertheless, our lipidomic and transcriptomic findings raise the possibility that CTL1-dependent choline availability influences the balance between lipid storage and utilization. How CTL1-dependennt choline transport is linked to mitochondrial metabolism remains unclear. CTL1 has been reported on the outer mitochondrial membrane (Michel & Bakovic, 2009), raising the possibility that altered choline availability following CTL1 loss could influence mitochondrial metabolism.

Another major finding of this study is that loss of *Ctl1* alters mTOR signaling. *Ctl1* SC-KO nerves showed increased mTORC1-associated S6 phosphorylation as well as increased mTORC2-associated AKT S473 phosphorylation. Increased mTORC1 activity has also been observed when choline metabolism is impaired in skeletal muscle (Taylor et al., 2017). Importantly, excessive mTORC1 activity in Schwann cells increases myelin abnormalities and causes hypermyelination (Beirowski, Wong, Babetto, & Milbrandt, 2017; Figlia, Norrmen, Pereira, Gerber, & Suter, 2017), while increased AKT signaling increases abnormalities in myelin architecture, including infoldings and outfoldings (Domenech-Estevez et al., 2016). mTORC2-mediated phosphorylation of AKT at the S473 promotes maximal AKT activation (Manning & Toker, 2017). Thus, increased mTORC1 and mTORC2-AKT signaling provides a potential mechanism contributing to the myelin abnormalities observed in *Ctl1* SC-KO nerves.

Interestingly, myelin infoldings and outfoldings are also characteristic of Charcot-Marie-Tooth disease variant 4B (CMT4B), which is caused by mutations in myotubularin-related proteins (Azzedine et al., 2003; Kiwaki et al., 2000; Othmane et al., 1999; Quattrone et al., 1996). Mouse models of CMT4B1 and CMT4B2 develop myelin outfoldings without alterations in myelin thickness (Bolino et al., 2004; Bolis et al., 2005; Bonneick et al., 2005; Robinson, Niesman, Beiswenger, & Dixon, 2008), resembling the phenotype observed in *Ctl1* SC-KO nerves. Notably, mTORC1 activity is elevated in CMT4B1 nerves (Guerrero-Valero et al., 2021).

The relationship between CTL1 loss or altered lipid metabolism and increased mTOR-AKT signaling is interesting. Cellular lipid and nutrient availability are closely integrated with mTOR signaling, and phosphatidylcholine synthesis influences membrane organization and the availability of lipid-derived signaling intermediates (He, Cho, & Blenis, 2025; Kenny et al., 2025). CTL1 deficiency could therefore affect mTOR activity indirectly through changes in cellular lipid composition. Conversely, enhanced mTOR signaling could itself influence lipid synthesis, storage, and catabolism. Our experiments do not establish the direction of this relationship. Determining whether normalization of mTOR signaling reduces the lipid and myelin abnormalities in *Ctl1* SC-KO nerves will help establish whether increased mTOR activity is a consequence of altered lipid homeostasis or contributes directly to the phenotype.

The effect of CTL1 loss also extends to the Schwann cell response to nerve injury. Choline and its derivatives have previously been implicated in remyelination in the CNS. For example, CDP-choline supplementation improves remyelination in an experimental multiple sclerosis model by increasing oligodendrocyte precursor proliferation and differentiation (Skripuletz et al., 2015). In Schwann cells, however, loss of CTL1 did not impair proliferation or the generation of repair Schwann cells after nerve injury. Remyelination of regenerated axons was also largely preserved, with no major changes in the number of myelinated axons or myelin thickness. Interestingly, altered mTOR signaling persisted during the injury response and later remyelination phase. Therefore, as during developmental myelination, Schwann cells can remyelinate axons in adult mice despite the loss of CTL1.

The relationship between choline metabolism, lipid homeostasis, and myelin structure may also have relevance to peripheral neuropathies. Lipid supplementation has recently been shown to ameliorate dysmyelination in peripheral neuropathies (Fledrich et al., 2018; Silva et al., 2025; Zhou et al., 2019). Choline derived lipids are reduced in several models of peripheral neuropathy including CMT1A and CMT1E (Kuipers et al., 2026; Silva et al., 2025). Schwann cell precursors derived from hiPSCs from CMT1A patients generate larger and more numerous lipid droplets, and dietary triglycerides supplementation improves myelination in a CMT1E model (Prior et al., 2024; Zhou et al., 2019). These findings raise the possibility that choline and lipid metabolism may represent modifiable pathways in peripheral neuropathies. Further studies will be required to determine whether choline supplementation or manipulation of related metabolic pathways can improve Schwann cell function in these disorders.

In conclusion, our findings show that CTL1 has a relatively minor role in regulating the overall extent of Schwann cell myelination but is important for maintaining normal myelin structure and lipid homeostasis. Loss of CTL1 alters myelin lipid composition, increases triglyceride accumulation, enhances mTORC1 and mTORC2-AKT signaling, and is associated with reduced expression of gene programs involved in fatty acid oxidation and oxidative phosphorylation. Despite these changes, Schwann cells remain capable of developmental myelination and remyelination after injury, revealing a notable difference between Schwann cells and oligodendrocytes in their dependence on CTL1.

### Methods and Materials Animals

All animal procedures were performed in accordance with the guidelines of the Institutional Animal Care and Use Committee (IACUC) at Rutgers University. Mice and rats were housed in microisolator cages on IVC racks in standard 12-hour light-dark cycles with food and water provided *ad libitum*.

*In vitro* fertilization of wildtype C57Bl6J (RRID:IMSR_JAX:000664) ova and C57/Bl6NJ (RRID:IMSR_JAX:005304) CTL1-targeted sperm (Slc44a1^[tm1a(EUCOMM)Hmgu]^; purchased from Baylor College, TX) was performed at the Rutgers University Genome Editing Core Facility to generate heterozygous CTL1^+/neo-LacZ^ mice. Upon generation of homozygous CTL1^neo-LacZ/neo-LacZ^ mice, they were crossed to FLPo mice (B6.129S4-*Gt(ROSA)26Sor^tm2(FLP*)Sor^*/J; RRID:IMSR_JAX:012930) to create a *Ctl1^flox/flox^* line with loxP sites surrounding exon 8. Subsequently these mice were crossed to Dhh^Cre+^ mice (FVB(Cg)-Tg(Dhh-cre)1Mejr/J; RRID:IMSR_JAX:012929) (Jaegle et al., 2003) to generate Dhh-Cre driven Schwann cell-specific *Ctl1* knockout mice (*Dhh-Cre^+^:Ctl1^flox/flox^)* and control mice (*Dhh-Cre^-^:Ctl1^flox/flox^* or *Dhh-Cre^+^:Ctl1^wt/wt^*). Experiments were conducted using age matched colony controls. Mice of either sex between 1-6 months of age were used for experiments. Sex differences were not examined in this study. Criteria for exclusion of animals during the experiment was animal illness or post-surgery illness. No animals or data were excluded during analysis. Surgeries for each cohort were performed on the same day and collection of tissue was performed in the same order as surgery to minimize confounders.

Timed pregnant SAS Sprague Dawley rats purchased from Charles River Laboratories were used for Schwann cell culture at P2 (detailed below). Rat embryos and pups are pooled regardless of sex.

### Surgical procedures

Sciatic nerve transection or crush injuries were performed on male or female mice aged 2-6 months as previously detailed (Patel et al., 2024). Briefly, under isoflurane, and in aseptic conditions, the sciatic nerve was exposed and transected or crushed distal to the sciatic notch. For crush injuries, the nerve was crushed with forceps for 30 seconds, then the forceps were rotated 90 degrees and the same site was crushed for an additional 30 seconds. The crush site was marked using charcoal coated forceps. The wound was sutured, and an analgesic was administered. The sciatic nerve was collected at different timepoints for experimentation.

### Cell cultures

HEK293FT (Invitrogen, Cat#: R70007; RRID:CVCL_6911) cells were used for lentiviral transduction (detailed below). Cells were grown in Dulbecco’s Modified Eagle Medium (DMEM) (Corning, Cat#: 10-017-CV), 10% fetal bovine serum (FBS) (Atlas Biologicals, Cat#: F-0500-D), with 1mM sodium pyruvate (ThermoFisher Scientific, Cat#: 11360070), 1X MEM Non-essential amino acids (ThermoFisher Scientific, Cat# 11140050), and Geneticin (G418 sulfate, Cat#: 11811031) at 37°C in a 10% CO_2_ incubator.

Primary rat Schwann cells were collected from P2 neonates as previously described (Monje, 2018). Briefly, sciatic nerves from P2 neonates were digested in 0.25% Trypsin (ThermoFisher Scientific, Cat#: 15050057) with 0.1% collagenase (Worthington Biochemical Corp. Cat#: LS004194). Afterward, nerves were centrifuged 50 x g for 5 minutes. The supernatant was removed, and the nerves were triturated with a glass pipette in DMEM with 10% FBS. The suspension was plated in media containing DMEM, 10% FBS, 1x Glutamax (ThermoFisher Scientific, Cat#: 35050079), and 1x Penicillin/Streptomycin (Corning, Cat#: 30-002-CI) onto 60mm plates coated with poly-L-lysine (PLL) (Millipore Sigma, Cat#: P7890) at a concentration of 8 sciatic nerves/60mm plate. Fibroblasts were removed by adding 10 µM Cytosine-ß-arabinofuranoside hydrochloride (Ara-C) to media for 3 days, followed by incubation with 20µg/ml mouse CD90 monoclonal antibody cloneT11D7e (Thy1.1) (Bio-Rad, Cat#: MCA04G) for 30 minutes followed by addition of 400µl rabbit serum complement (Millipore-Sigma, Cat# 234400). Cells were then expanded and grown in Schwann cell growth media: DMEM, 10% FBS, 1x Glutamax, 1x Penicillin/Streptomycin, 10 ng/ml Recombinant Human NRG1-beta 1/HRG1-beta 1 EGF domain (Nrg1-EGF domain) (VWR, Cat#: 10025-698), and 2µM forskolin (Millipore-Sigma, Cat# F3917) for experimentation.

Primary DRGs were collected from E15 rat embryos as previously described (Monje, 2018). Briefly, E15 rats were dissected, and their spinal cords exposed. DRGs were plucked from spinal cords and dissociated with 0.25% trypsin and plated at a density of 1.35 DRG/coverslip on coverslips precoated with growth factor reduced Matrigel (Corning, Cat#: 356231). DRGs were grown in DRG culture media: neurobasal media (ThermoFisher, Cat#: 21103-049), with 0.08% D-(+)-glucose (Millipore-Sigma, Cat#: G27528), 1x penicillin/streptomycin, 1x B-27 supplement (ThermoFisher Scientific, Cat#: 17504-044), 1x Glutamax, and 50 ng/ml 2.5S nerve growth factor (NGF) (Fisher Scientific, Cat#: NC0419584). To get a pure neuron culture, media was supplemented with 10µM 5-Fluoro-2′-deoxyuridine (FUDR) and 10µM Uridine (U) for 3 days. Afterward, DRG were maintained in DRG culture media with media changed every 2 days.

### Lentiviral Transduction

Lentiviral transduction was performed as previously described (Heffernan et al., 2017). Briefly, a 21-nucleotide shRNA (gatacgacagctatggaaata) was generated to target *Ctl1* at positions 192-213. The sequence was cloned into a pLL3.7 vector (Addgene, Cat#: 11795) (Rubinson et al., 2003).

HEK293FT cells were passaged onto 100mm cell culture plates coated with PLL; 5 × 10^6^ cells/plate. The next morning, cell media was changed to Advanced DMEM (Thermo-Fisher Scientific, Cat#: 12491051) with 2% FBS, 1x Glutamax, 1% chemically defined (CD) lipid concentrate, and 0.03mM cholesterol (Mllipore-Sigma, Cat#: C4951). Cells were transfected using a calcium phosphate transfection kit (Thermo-Fisher Scientific, Cat#: K278001) using 11µg packaging plasmid psPAX2 (Addgene, Cat#: 12260) and 5.5µg envelope plasmid PMD2.G (Addgene, Cat#: 12259) with 16.5µg sh*Ctl1* pLL3.7, and 0.25M CaCl_2_. After 48 hours, viral supernatant was collected and mixed in a 3:1 ratio with lenti-X-concentrator (Takara, Cat#: 631232) and added to Schwann cell cultures in Schwann cell growth media. After 24 hours, media was changed to Schwann cell growth media.

### RT-PCR and qPCR

RNA from rat cell cultures and tissue was extracted with TRI reagent (Sigma, catalog#: T9424), and cDNA was generated with the SuperScript III first-strand synthesis kit (Invitrogen, catalog#: 18080051). *Ctl*-family cDNA was amplified using the following primers: *Ctl1*: Fwd 5’-gcaatagcgaacagtggcc, Rev 5’-acttctggacgtcactcagg; *Ctl2*: Fwd 5’-ggacgcctcagaaatatgacc, Rev 5’-cctacgatggccaggaagag; *Ctl3*: Fwd 5’-actcagagtccaagtgcaga, Rev 5’-ggtgtctcttcctgccatga; *Ctl4*: Fwd 5’-ggcaaccccagtacgtctat, Rev 5’-tgaagagagaacgttgggct; *Ctl5*: Fwd 5’-tccatgatgcttcgcctaca, Rev 5’-actgccgatagtattcccagt.

For mouse sciatic nerve qPCR, RNA was extracted as described below (see Collection of nerves for RNA-seq analysis) and cDNA was generated with SuperScript III. The following primers were used: *Ctl1* exon 7-8: Fwd 5’-agttctggtcatactgggttca, Rev 5’-cactgtcgctgaaatggcat; *Ctl1* exon 3-4: Fwd 5’-ggaagcaataccgaacagtgg, Rev 5’- tgcaaacttctggacgtcac; *GAPDH*: Fwd 5’-ccccatgtttgtgatgggtg, Rev 5’-ggcatggactgtggtcatgag. qPCR was performed on a LightCycler 480 II (Roche Applied Science) using the Maxima SYBR Green/ROX qPCR master mix (Thermo Fisher Scientific, catalog#: K0222).

### Collection of nerves for RNA-seq analysis

Mice were euthanized with CO_2_ and both intact sciatic nerves were collected per animal. Nerves were placed into ice cold DEPC treated PBS with RNasin where the epineurium was removed from intact nerves and then nerves were snap frozen in liquid nitrogen. Nerves were then placed into a 2ml bead mill tube with 1ml Tri-Reagent and homogenized in a Fisherbrand Bead Mill 24 homogenizer for 30sec at 5m/s for 2 cycles. Chloroform was added and mixed and the tubes were centrifuged. The aqueous phase was removed and mixed with 70% EtOH. The mixture was then centrifuged through the NucleoSpin RNA Mini Kit (Macherey-Nagel, Cat#: 740955.50) and RNA extraction was completed following the manufacturer instructions. RNA was dissolved in RNAse free water and samples were sent to PrimBio Research Institute for mRNA enrichment, library preparation, sequencing, and alignment. Raw reads from PrimBio were analyzed with R version 4.5.1 in R Studio using DEseq2 to obtain differential expression and normalized counts data (Love, Huber, & Anders, 2014).

Gene set enrichment analysis was performed using GSEA software (Mootha et al., 2003; Subramanian et al., 2005). Volcano plots were made with Enhanced Volcano R package (Blighe K, 2024).

### Transmission electron microscopy, morphometric analysis, and toluidine blue staining

Nerves for electron microscopy and toluidine blue staining were fixed in 4% PFA (Electron Microscopy Sciences, Cat#: 15710) and 2.5% glutaraldehyde (Electron Microscopy Sciences, Cat#: 16365) in 0.1M sodium cacodylate (Electron Microscopy Sciences, Cat#: 12300) in phosphate buffer (Electron Microscopy Sciences, Cat#: 19340-72) for 24 hours at 4°C. Nerves were postfixed with 1% OsO_4_ (Electron Microscopy Sciences, Cat#: 19152) and 1.5% KFeCN (Electron Microscopy Sciences, Cat#: 25154) in phosphate buffer for 5 hours then washed with distilled water. Nerves were then dehydrated with graded ethanol steps. Nerves were then infiltrated with propylene oxide (Electron Microscopy Sciences, Cat#: 20412) followed by gradual replacement with resin: Embed 812 (Electron Microscopy Sciences, Cat#: 14900), DDSA (Electron Microscopy Sciences, Cat#: 13710), NMA (Electron Microscopy Sciences, Cat#: 19000) and DMP-30 (Electron Microscopy Sciences, Cat#: 13600) in a 2:1.6:0.8:0.077ml ratio. Nerves were embedded in resin at 65°C for 24 hours. For EM, nerves embedded in resin were sent for sectioning and processing to the Rutgers Robert Wood Johnson Medical School Core Imaging Lab. Sections were imaged using a ThermoFisher-FEI Tecnai12 Bio-twin TEM. Analyses were performed using Fiji (Schindelin et al., 2012).

*G*-ratios were calculated using MyelTracer (Kaiser et al., 2021). Fibers possessing myelin abnormalities were excluded from the quantification. At least 100 axon-myelin units were counted per individual animal.

For semi-thin toluidine sections, blocks were sectioned using a Leica EM UC7 ultramicrotome at a 300-500nm thickness. Sections were collected onto glass coverslips and stained with a 1% toluidine blue (Millipore Sigma, Cat#: T3260), 1% sodium borate solution in distilled water for 5-10 minutes. Excess stain was washed away, and coverslips were mounted with Permount (Fisher Scientific, Cat#: Sp15-100). Images were taken with a Hamamatsu Orca-ER camera using a 60x objective on a Nikon Eclipse TE2000-U Epi-fluorescence microscope. Whole nerve cross sections were imaged and counted for myelin abnormalities in intact nerves.

### Endogenous Choline Assay

Intracellular choline was measured as described previously (Heffernan et al., 2017). Briefly, control and *Ctl1* knockdown Schwann cells, at a 2-3x10^5^ cell density, were collected, cellular ions (H^+^/Na^+^/K^+^) were exchanged for Cs^+^ with ice-cold 100mM cesium chloride, resuspended in methanol, and sonicated. Methanol suspended lipids samples were mixed with 50mM deuterated choline standard (d9, Cambridge Isotopes Labs, Cat#: DLM-5491) and spotted to MALDI-TOF plates. Assessment of intracellular choline (m/z 104) and d9-choline (m/z 113) was made on a 4800 Plus MALDI TOF/TOF analyzed.

### Whole nerve lipidomics

For whole sciatic nerve lipidomics, mice were euthanized with CO_2_ and both intact sciatic nerves were collected per animal. Nerves were placed into ice cold PBS and the epineurium was removed and then nerves were snap frozen in liquid nitrogen. 0.5mL of an extraction solvent of 0.1M hydrochloric acid, 4.95mL methanol, and 50µL of SPLASH lipidoMIX Internal Standard (Avanti Polar Lipids 330707) was added and nerves were homogenized in a bead mill homogenizer for 30s at 5m/s for 2 cycles. Then, 1mL of methyl tert-butyl ether (MTBE) (Sigma, catalog#: 34875) was added and the tube was vortexed for 30s. 600µL of the top MTBE layer was transferred to a new tube and dried under nitrogen gas. Samples were sent to the Rutgers Metabolomics Shared Resource for LC-MS analysis.

### Myelin Fractionation Lipidomics

Myelin was fractioned from whole sciatic nerve extract (Erwig et al., 2019). Dissected sciatic nerves from control and *Ctl1* SC-KO P30 mice were snap frozen in liquid nitrogen and ground in a chilled ceramic mortar. The nerve powder was collected, suspended in cooled 1ml of 0.32M sucrose in 10 mM Tris buffer, pH 7.4 containing cOmplete mini protease inhibitor cocktail (Millipore Sigma, Cat#: 11836170001) and homogenized with a Dounce homogenizer. The homogenate was filtered through a 200μm nylon mesh (Spectrum Labs cat# 145811) to remove the bulk of the collagen and carefully layered over 4ml of 0.9M sucrose in 10 mM Tris buffer, pH 7.4, in a Nalgene spin tube (Thermo, catalog#: 3118-0050) and centrifuged at high speed (75,000 x *g*) in a Beckman Coulter Optima L-100XP Ultracentrifuge, Rotor SW55Ti for 2 hours. 200µL of the myelin fraction at the 0.32M/0.9M sucrose interface was collected, osmotically shocked in cold 800µL 20mM Tris·HCl, and homogenized five to seven strokes in a Dounce homogenizer. Myelin was pelleted and washed by successive centrifugation and wash steps in 20mM Tris·HCl, 15 mins at 4°C (at 75,000 x *g* and 12,000 x *g*; rotor SW55Ti). Wet weight of the final myelin pellets were established and subjected to lipidomic LC-MS/MS analysis at the Washington University Metabolomics Facility.

### Immunofluorescence

Mice were euthanized under CO_2_ and nerves were dissected out and fixed in 4% PFA in PBS solution for 1 hour. After washing, nerves were cryopreserved in 30% sucrose in water for 24-48 hours. Nerves were then embedded using tissue freezing media (General Data, Cat#: TFM-(color)) and frozen at -20°C. Nerves were then sectioned using a Leica CM3050S cryostat at a 12µm thickness for all sections.

For staining tissue, nerve sections were thawed at room temperature for 10 minutes then rehydrated in PBS. Sections were blocked and permeabilized in 0.3% Triton X-100, 0.1M glycine, with 5% normal donkey serum for 1 hour. Primary antibodies were diluted into PBST with 5% normal donkey serum and sections were placed at 4°C overnight. The next day, sections were washed with PBST and then secondary antibody diluted in 0.3% Triton X-100 with 5% normal donkey serum added for 1 hour at room temperature. After washing, DAPI (4’,6-diamidino-2-phenylindole) (Fisher Scientific, Cat#: D1306, 5 μg/mL) was added for 10 minutes in PBS and slides were mounted with Vectashield Vibrance antifade mounting media (Vector Laboratories, Cat#: H-1700-10).

Primary antibodies used for immunofluorescence: Ki67 (Cell Signaling Technology, Cat#: 9129, RRID: AB_2687446, 1:400), c-JUN (Cell Signaling Technology, Cat#: 9165S, RRID: AB_2130165, 1:800), EGR2 (KROX20, Abcam, Cat#: AB245228, RRID: AB_2934181, 1:1000), SOX10 (Fisher Scientific, Cat#: AF2864, RRID: AB_442208, 1:150), pS6 S235/S236 (Cell Signaling Technology, Cat#: 4858S, RRID:AB_916156, 1:500), and pAKT S473 (Cell Signaling Technology, Cat#: 4051S, RRID:AB_331158, 1:500).

Secondary antibodies used for immunofluorescence: donkey anti-rabbit Alexa Fluor 647 (Jackson ImmunoResearch, Cat#: 711-605-152, RRID:AB_2492288), donkey anti-goat Alexa Fluor 488 (Jackson ImmunoResearch, Cat#: 705-545-003, RRID:AB_2340428), donkey anti-rabbit Rhodamine (TRITC) (Jackson ImmunoResearch, Cat#: 711-025-152, RRID:AB_2340588), donkey anti-rabbit Alexa Fluor 488 (Jackson ImmunoResearch, Cat#: 711-545-152, RRID:AB_2313584), and donkey anti-rabbit Alexa Fluor 488 (Jackson ImmunoResearch, Cat#: 711-545-152, RRID:AB_2313584). All secondary antibodies were used at a 1:500 dilution.

All images were taken with a Hamamatsu Orca-ER camera using a 20X or 40X objective on a Nikon Eclipse TE2000-U Epi-fluorescence microscope. For nerve sections, 3-5 images were taken per nerve section. Image analysis was performed using Fiji.

### Western blot analysis

After dissection, sciatic nerves were frozen with liquid nitrogen. Nerves were homogenized in bead mill tubes with lysis buffer containing 150mM Tris-HCl (pH6.8), 6% sodium dodecyl sulfate (SDS), 0.001% bromophenol blue, 10% B-mercaptoethanol, 50mM NaF, 1mM NaVO_4_, and cOmplete mini protease inhibitor cocktail (Millipore Sigma, Cat#: 11836170001) in a Fisherbrand Bead Mill 24 homogenizer with 2 30-second cycles at 5m/s. Homogenized nerves were kept on ice for 15 minutes, heated to 95°C for 3 minutes, and centrifuged 16,200*g* for 15 minutes at 4°C. Supernatant was removed and stored at -80°C.

Lysates were loaded onto SDS-PAGE gels and then transferred to PVDF membranes. Membranes were stained for total protein with Ponceau S (Fisher Scientific, Cat#: BP103-10). Membranes were blocked using 2% blotting-grade blocker (Biorad, Cat#: 1706404) in TBS for 1 hour. Membranes were incubated with primary antibody in 5% BSA in TBST overnight at 4°C. Membranes were washed in TBST, then incubated with secondary antibodies diluted in 2% blotting-grade blocker in TBST for 1 hour, followed by washing with TBST. Membranes were imaged using a BioRad ChemiDoc MP imaging system.

Primary antibodies used for Western blotting: MPZ (Millipore Sigma, Cat#: AB9352, RRID: AB_571090, 1:500), MBP (rabbit, EMD Millipore, Cat#: AB980, RRID: AB_92396, 1:1000, Calnexin (Enzo Life Sciences, Cat#: ADI-SPA-860-D, RRID: AB_2038898, 1:1000), pS6 S235/S236 (Cell Signaling Technology, Cat#: 4858S, RRID:AB_916156, 1:2000), tS6 (Cell Signaling Technology, Cat#: 2317S, RRID:AB_2238583, 1:1000), pAKT S473 (Cell Signaling Technology, Cat#: 4051S, RRID:AB_331158, 1:1000), pAKT T308 (Cell Signaling Technology, Cat#: 2965S, RRID:AB_2255933, 1:1000), and tAKT (Cell Signaling Technology, Cat#: 9272S, RRID:AB_329827, 1:1000).

Secondary antibodies used for Western blotting: goat anti-rabbit Alexa Fluor 680 (Jackson ImmunoResearch, Cat#: 111-625-144, RRID:AB_2338085), donkey anti-mouse Alexa Fluor 790 (Jackson ImmunoResearch, Cat#: 715-655-150, RRID:AB_2340870), donkey anti-rabbit Alexa Fluor 647 (Jackson ImmunoResearch, Cat#: 711-605-152, RRID:AB_2492288), donkey anti-rabbit Rhodamine (TRITC) (Jackson ImmunoResearch, Cat#: 711-025-152, RRID:AB_2340588), donkey anti-rabbit Alexa Fluor 488 (Jackson ImmunoResearch, Cat#: 711-545-152, RRID:AB_2313584) and donkey anti-chicken Alexa Fluor 790 (Jackson ImmunoResearch, Cat#: 703-655-155, RRID:AB_2340382). All secondary antibodies were used at a 1:10000 dilution.

### Experimental Design and Statistical Analysis

Experimenter blinding was performed for all the analyses. Sample sizes of 3 to 5 animals per experimental condition were used, similar to the range used in previously published studies in the field (Arthur-Farraj et al., 2012; Beirowski et al., 2017; Daboussi et al., 2023; Heffernan et al., 2017; Li, Banton, Min, Parkinson, & Dun, 2021; Napoli et al., 2012; Norrmen et al., 2018; Reed et al., 2020; Roberts et al., 2017). Statistical analysis was performed using GraphPad Prism software version 10.1.0. The following tests were used: Two-tailed Paired t-test, One-tailed Unpaired t-test, Two-tailed Unpaired t-test with Welch’s correction, One sample t and Wilcoxon test, and Two-way ANOVA with Uncorrected Fisher’s LSD. Statistic test used for individual experiments are listed within Figure Legends. Data are presented as Mean ± SEM, where p ≤ 0.05 was considered significant. P-values are displayed in graphs.

## Acknowledgments

We thank the Rutgers Robert Wood Johnson Medical School Core Imaging Laboratory for assistance with electron microscopy and the Rutgers Metabolomics Shared Resource and Washington University School of Medicine Metabolomics Facility for lipidomic analyses. This work was supported by the NIH-NINDS R01NS118020 and R21NS140995 to H.A.K.

